# Emergent vulnerability to intensive coastal anthropogenic disturbances in mangrove forests

**DOI:** 10.1101/2021.04.17.440255

**Authors:** Yangfan Li, Zhen Zhang, Yi Yang, Yi Li

## Abstract

Mangrove forests, as one of the most productive coastal ecosystems in tropical and subtropical areas, provide multiple valuable ecosystem services for human well-being. Mangrove coverage has been declining dramatically across much of developing regions due to extensive coastal anthropogenic disturbances such as reclamation, aquaculture, and seawall construction. As coastal human activities increase, there is urgent need to understand not only the direct loss, but also the vulnerability of mangroves to anthropogenic disturbances. In this study, we evaluated spatial pattern of mangrove vulnerability based on the conceptual framework of “Exposure-Sensitivity-Resilience” using geospatial datasets in mainland China. We find that within all 25,829 ha mangroves in five coastal provinces of mainland China in 2015, nearly 76% of mangroves was exposed or threatened by anthropogenic disturbances. Coastal reclamation and aquaculture were the key threats causing mangrove vulnerability. The overall distribution of high, medium and low vulnerability was following similar trend of aquaculture distribution, which suggests aquaculture was the greatest anthropogenic disturbance agent to mangroves. Hotspot regions for mangrove vulnerability are located at the developing provinces such as Guangxi and Hainan. This study provides the first spatially explicit evidence of the vulnerability of mangrove forests to intensive coastal anthropogenic disturbances at national scale, cloud serve as a benchmark for navigating coastal ecological redline management and coastal ecosystem restoration.

## 1 Introduction

Global assessments indicate that coastal tropical and subtropical areas are threatened by the greatest urbanization rates, suggesting an alarming to the conservation of coastal ecosystems in this regions and their associated ecosystem services (Branoff, 2017; Friess et al., 2016; Rocha et al., 2018; Seto et al., 2012). Mangrove provides essential ecosystem services to coastal community, including products provision, cultural services, carbon sequestration, water purification, shoreline protection/regulation and biodiversity conservation (Bell & Lovelock, 2013; Crooks et al., 2018; Estoque et al., 2018; Friess & Webb, 2014). However, intensive anthropogenic disturbances caused by rapid coastal development exacerbates degradation of coastal systems. Critical functions of coastal wetland ecosystems have been eroded by global-scale anthropogenic pressures, which leads to irreversible land use and cover change, massive biodiversity loss and overloaded land-based sources of pollution (Duke et al., 2014; Wang et al., 2018).The world’s area of mangrove forests has decreased by about 35% globally from 1980 to 2000 and experienced a decline of 1.97 % annually from 2000 to 2012 (Hamilton & Casey, 2016; Valiela et al., 2001), about 3-5 times greater than that of terrestrial forest loss (Duke et al., 2014). Coastal reclamation and conversion of mangroves to aquaculture or agriculture creates pressure on mangrove ecosystems and has thus been a major driver of mangrove destruction (Flores-de-Santiago et al., 2017; Mukherjee et al., 2014; Spalding, 2010). Numerous studies have quantified increasing coastal reclamation and modifications over time, with demonstrated the deleterious effects of urban, agricultural, aquaculture, and infrastructure development on mangrove ecosystem (Hamilton & Friess, 2018; Rivera-Monroy et al., 2017). Global sustainability science increasingly recognize the changes of structure and function taking place in ecosystem, understanding which ecological functions are vulnerable to anthropogenic disturbances is key to sustainable development (Richards & Friess, 2016).

Vulnerability defines the degree to which a system or system component is susceptible to experience harm due to exposure to a perturbation or stressor which mostly associated with environmental, social change, and from the absence of capacity to adapt (Adger, 2006; Turner II et al., 2003; White, 1974). Vulnerability analysis in coupled social-ecological systems draws on three major concepts: exposure, sensitivity and resilience (IPCC, 2014; Turner II et al., 2003). Through many years of studies on mangroves under climate change to reveal its distinct structures and process (Lovelock et al., 2015; Duke et al., 2014; Webb et al., 2013; Xu et al., 2016), there remains limited conceptual framework or model for the vulnerability of mangroves to anthropogenic disturbances (Branoff, 2017; Mukherjee et al., 2014; Ventura & Lana, 2014). Predictions of mangrove vulnerability around cities are mostly reported in low- and lower-middle-income regions of Africa, Latin America and India (DasGupta & Shaw, 2013; Elmqvist et al., 2013; Nortey et al., 2016).

Urbanization, in terms of landscape pattern changes, is particularly detrimental to degradation of mangrove in the Anthropocene. Human-induced coastal reclamation, as a typical land use change, is one of the key drivers of mangrove deforestation (Richards & Friess, 2016). Landscape changes caused by the human activities of coastal ecosystems can produce both direct and indirect effects on the long term sustainability of mangrove wetlands and coastal communities (Koh & Khim, 2014; Yim et al., 2018). These changes raise questions such as: how to identify the pattern of vulnerable mangroves to multiple environmental changes, and what are principal threats to local mangroves? This recognition requires revisions and enlargements in the basic design of vulnerability assessments, including the capacity to treat coupled social-ecological systems and those linkages within and without the systems that affect their vulnerability.

Here, our aim was to evaluate the spatial pattern of mangrove vulnerability to coastal anthropogenic disturbances in mainland China, the regions where experienced a rapid and unprecedented process of urbanization. Our specific research questions were: (1) what is the spatial pattern of mangrove distribution across mainland China? (2) what is the spatial pattern of mangrove vulnerability to reclamation in mainland China? (3) what are the principle disturbance agents to mangroves? For addressing these three questions, we firstly mapped the mangrove extent for mainland China by using high-resolution Google Earth images and visual interpretation, and subsequently developed a methodology to evaluate the vulnerability of mangrove forests to three coastal anthropogenic disturbances (i.e., reclamation, aquaculture, and seawall).

## 2 Study area

Since half of global mangrove areas are distributed within 25 km of urban centers inhabited by dense human population (McLeod & Salm, 2006; Millennium Ecosystem Assessment, 2005), more than 90% of the world’s mangroves are destroyed or threatened by diverse forms of human activities in recent decades (Mcleod et al., 2011; Murray et al., 2018; Silliman et al., 2009). Alarming losses of mangrove cover have occurred in most of developing areas of China due to extensive coastal reclamation, deforestation, engineering and urbanization. Although mangroves in China cover only 0.14% of the global mangrove area, one third of mangrove species can be found in this region (Romañach et al., 2018), suggesting that valuable contribution of mangrove conservation in China to biodiversity conservation in global scale.

Natural mangroves grow along the southeast coast from Fujian Province to Hainan Province (18°12’-27°20’N). Planted mangroves are scattered within boundary of natural mangroves. To protect local mangrove ecosystems, there are 35 conservation areas and several Ramsar Convention sites in China, e.g., Dongzhaigang Mangrove Nature Reserve in Hainan Province, Fujian Zhangjiangkou National Mangrove Nature Reserve.

Extensive coastal reclamation occurred in rapid urbanizing metropolitans of mainland China, such as Bohai bay, Yangtze River delta and Pearl River delta, was responsible for approximately 950,000 ha coastal wetlands loss and consequently ecosystem service decrease in recent years (Sajjad et al., 2018; Tian et al., 2016). According to the statistical data from the State Oceanic Administration, People’s republic of China in 2015, the impacts of coastal reclamation on ecosystem services resulted in the loss in wetlands of about $31,000 million, which accounts for 6% of ocean economy in China (State Oceanic Administration, 2016).

## 3 Materials and Methods

### 3.1 Mangrove mapping

For assessing the spatial vulnerability of mangroves, we firstly mapped the extent of mangrove forest in mainland China using artificial visual interpretation. Processes of mangrove interpretation included extracting mangroves with difference sources of base map (global mangrove datasets, Landsat images, etc.) and modification with filed verification in each province. Prior to interpretation, we imported the global mangrove distribution vector data in 2000 (https://www.usgs.gov/) as a base map and selected some mangrove samples in all five provinces as test examples for mangrove identification. By modifying these samples under the guidance of mangrove experts, mangroves are shown as dark green ribbon pattern and uniform texture feature along coastline (Fig. S1). To eliminate visual interference from salt marsh species (e.g., *Spartina alterniflora*), we selected Google Earth images in winter 2015 because mangroves can be more easily identified during dormant season of salt marshes (Fig. S2). Meanwhile, considering tidal inundation to mangroves, we compared the images of different time period in same areas and selected images with low tide. This procedure helped to eliminate the error in extracting submerged mangroves. Ultimately, we saved the mangrove layer in Google Earth and exported to ArcMap 10.2. After interpretation with remote sensing images, we verified mangrove area by field survey from February 2016 to March 2018 to examine interpretation accuracy of mangrove forest map. We selected 56 mangrove validation sites and identified mangrove distribution using Unmanned Aerial Vehicle (UAV) vertical photography and field survey. About 758 UAV images were used to create a confusion matrix indicating the producer’s and user’s accuracy. Availability of aerial photographs in our study area facilitated ground truth process in evaluating accuracy of classified mangroves.

### 3.2 Detection of anthropogenic disturbances

#### 3.2.1 Reclamation activities

Reclamation is defined as the conversion of coastal land to agricultural, industrial, and urban land use (Tian et al., 2016). We assume that the impacts of reclamation activities are decreasing with distance to artificial coastline, which means the longer distance to coastline the less negative influences of reclamation activities to local mangroves. We created a buffer of coastline sourced from Sajjad et al., 2018, and the buffer radius is set as 900 m since this distance covers 98% of mangroves in mainland China. Then we conduct a stress-gradient analysis in three levels of pressure based on 300 m intervals (Fig. 1, adapted from Sutton-Grier et al., 2015). The coastal area within the first 300 m buffer zone adjacent to reclamation suggests the highest level of pressure from reclamation activities on mangroves, which was assigned with a high-pressure value.

**Fig. 1.**
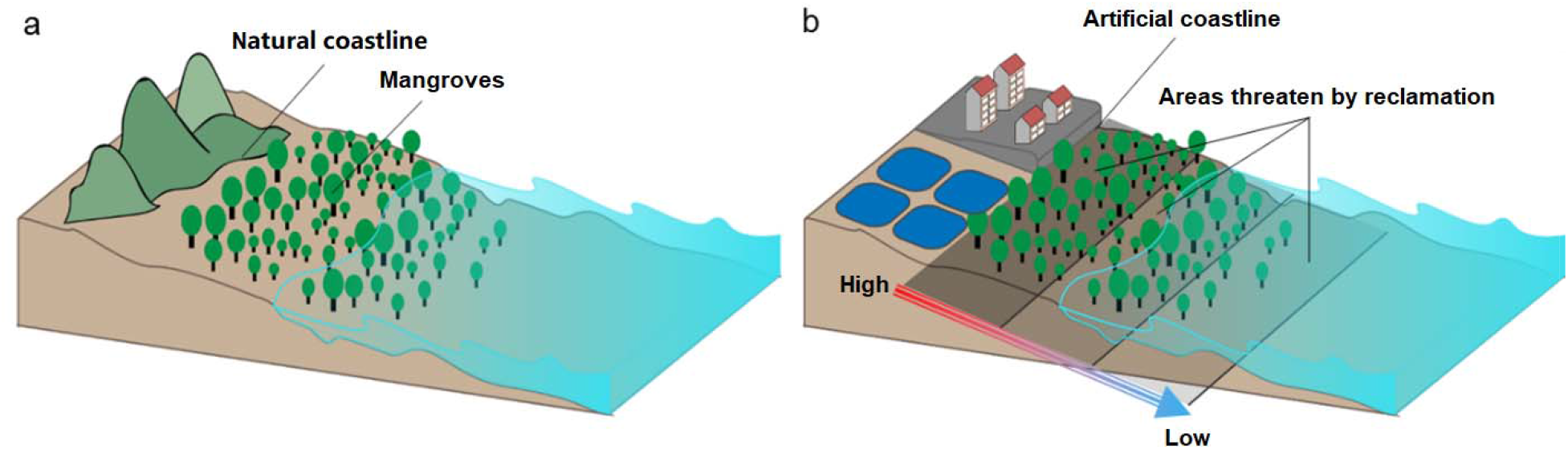
A schematic diagram of the spatial distribution of the reclamation pressure.

#### 3.2.2 Aquaculture pollution

Aquaculture pollution dataset contains two layers-aquaculture ponds and tidal creek, which are the main sources of pollutants in local ecosystem. Muddy sediments and wastewater of aquaculture ponds produce highly concentrated pollutants that lead to degradation of mangrove system (Xin et al., 2014). Aquaculture ponds data was extracted from the layer of coastline in our previous study (Sajjad et al., 2018), tidal creek was detected by visual image interpretation based on Google Earth. We created buffer zones for both layers with distance of 70 m, 140 m and 210 m separately to model stress-gradient analysis regarding aquaculture pollution (Fig. 2, adapted from Sutton-Grier et al., 2015). High threats from aquaculture pollution was assigned in the coastal area of 70 m buffer zone, while relative low threats within the 210 m buffer zone.

**Fig. 2.**
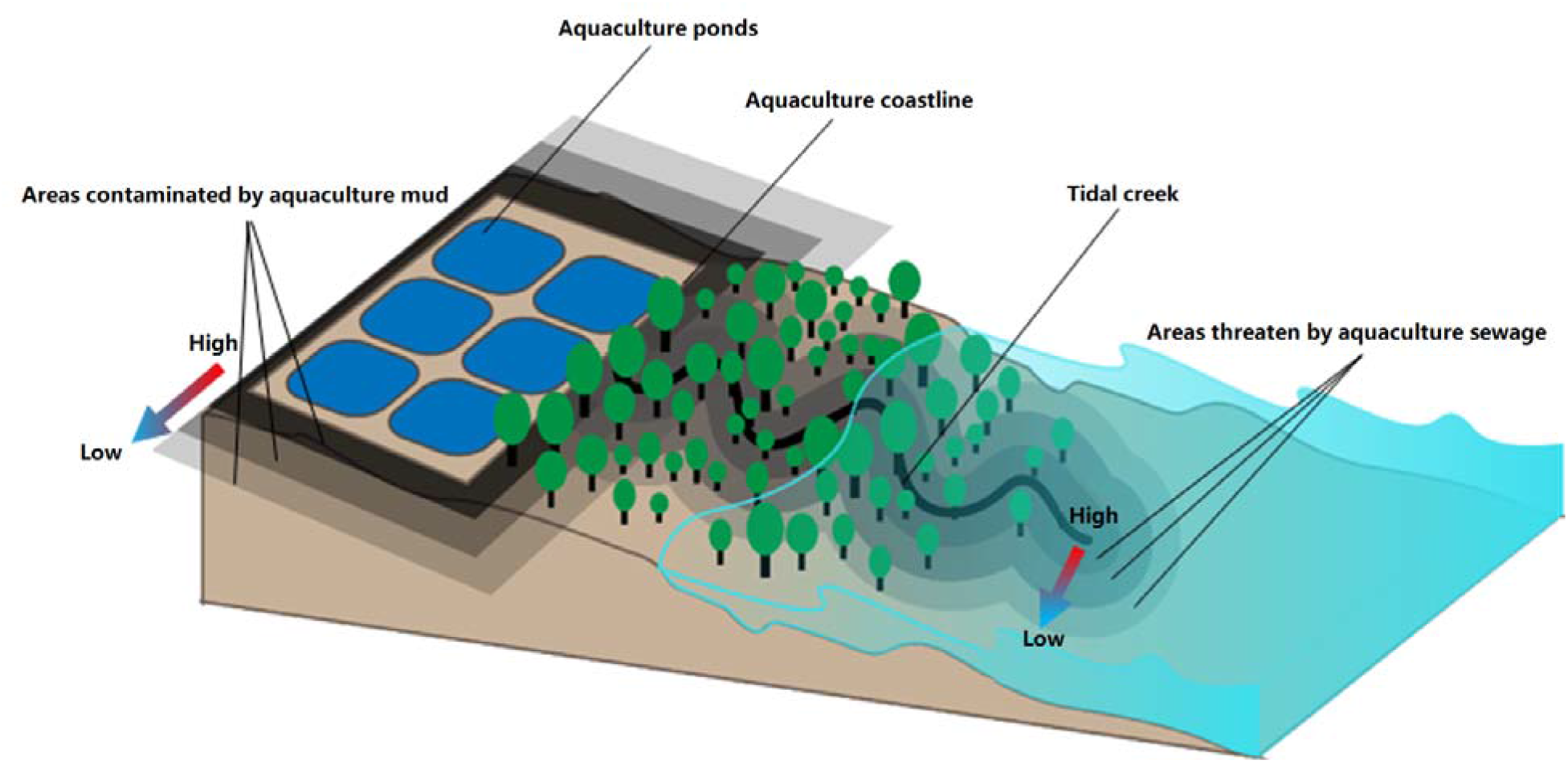
A schematic diagram of the spatial distribution of the aquaculture pollution pressure.

#### 3.2.3 Seawall

With the increase of dam construction and large reservoir projects in developing countries (Moran et al., 2018; Shi et al., 2018), large area of landscape transformed from natural coastline to seawall. The newly built impervious surface areas squeeze coastal zone toward ocean with increasing risk of flood (Doody, 2013; Ramesh et al., 2015). Three buffer zones were created along the claimed coastline with the distances of 300 m, 600 m and 900 m to create stress-gradient layer of flood risk (Fig. 3, adapted from Sutton-Grier et al., 2015). In this case, 300 m buffer zone is the area with the lowest flood threat, which is opposite to the threat distribution of reclamation threat.

**Fig. 3.**
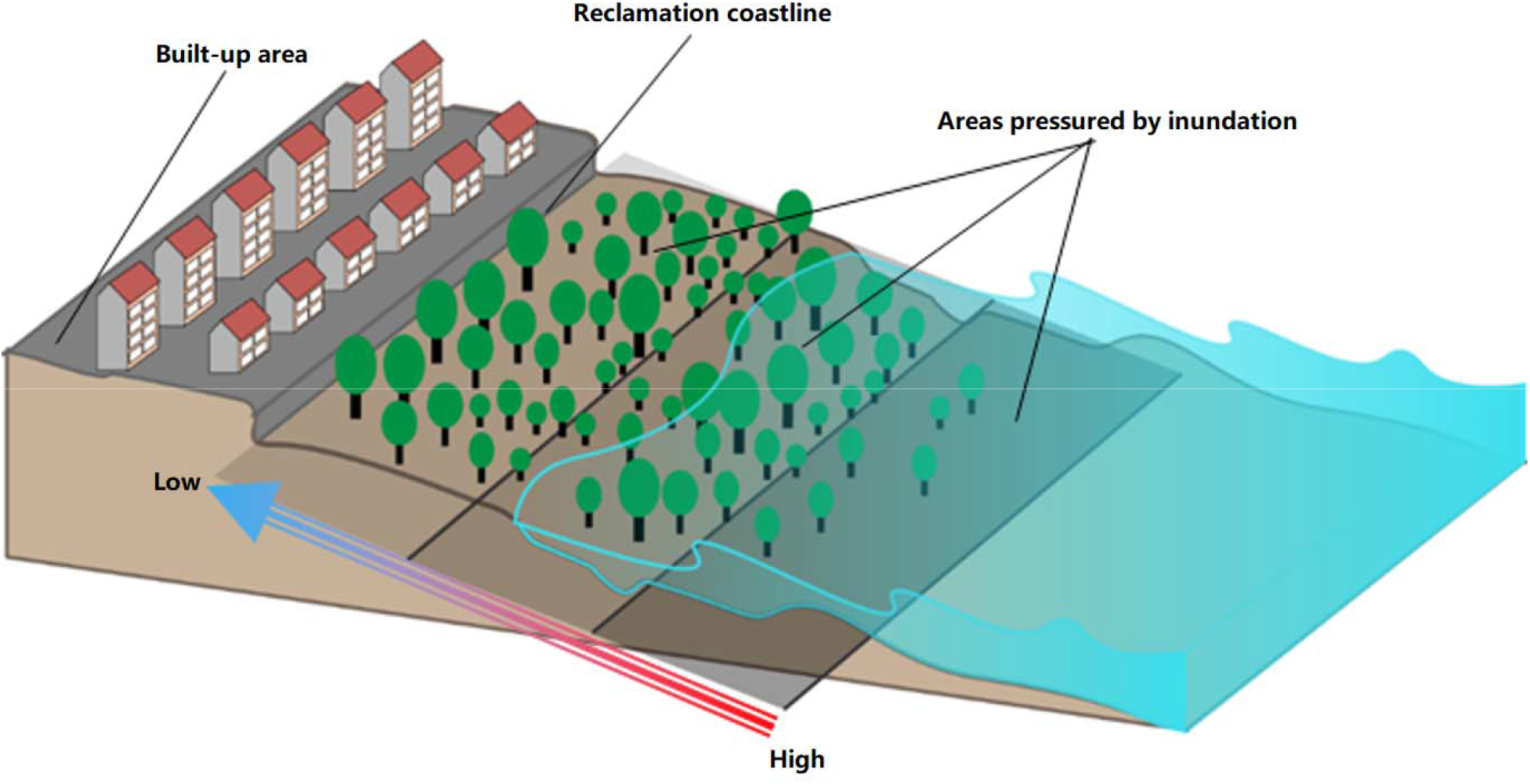
A schematic diagram of the spatial distribution of the seawall barrier pressure.

### 3.3 Vulnerability assessment

Referring to the vulnerability framework proposed by Turner II et al. (2003), we proposed a spatial assessment framework to evaluate mangrove vulnerability using InVEST model, which is a spatially-explicit modelling tool for assessing ecosystem services and would return a suite of results in a raster format to evaluate ecosystem states. (Sharp et al., 2015). We used the Habitat Risk Assessment submodule from InVEST model with inputs of all anthropogenic disturbance agents to local mangroves. Vulnerability of mangroves was characterized by integrating three subsystems: exposure, sensitivity and resilience (Fig. 4; Table S1).

**Fig. 4.**
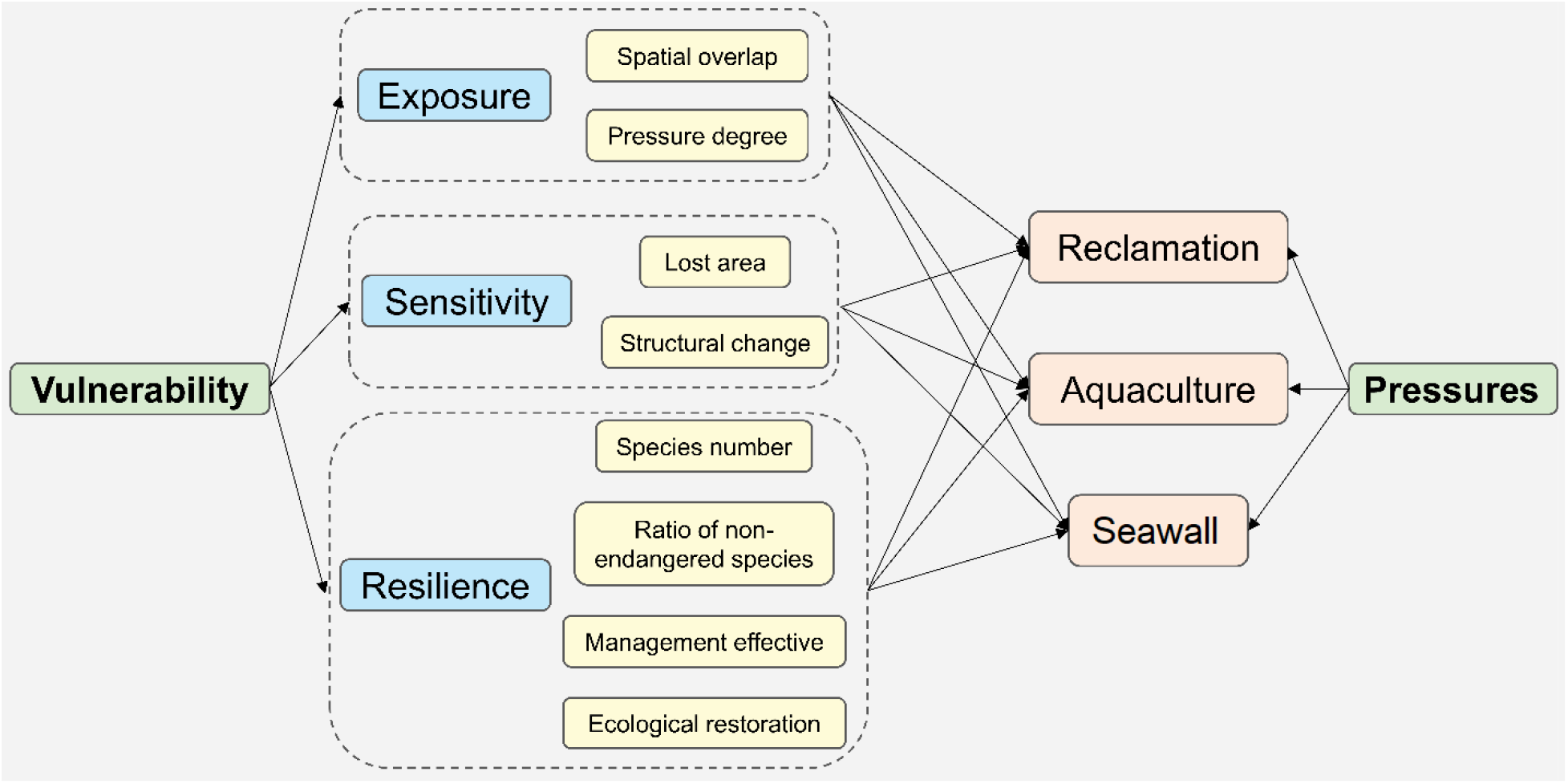
Indexes of mangrove vulnerability to anthropogenic disturbance pressures.

Exposure index represents the amount and intensity of anthropogenic stressors to mangroves. Exposure value was divided by the overlapped area of mangroves and reclamation and reclamation pressure intensity. Sensitivity index indicates negative impacts of reclamation on mangroves, and was quantified by mangrove loss area and mangrove species change. Resilience characterizes the ability of mangroves to maintain their structural and functional stability in the process of and after reclamation disturbances. And it was quantified by mangrove species number, ratio of non-endangered species, regional ecological management, and potential restoration areas for mangroves. We then clustered these indexes to corresponding vulnerability ranking of low (value=1), moderate (value=2) and high (value=3) using *K*-means algorithm (See Supplementary material for more details).

Exposure indicators represent risks of mangrove while experiencing vast reclamation activities, which describes characteristics and components of exposure to reclamation. Inputs of exposure included polygons of mangroves and reclamation with attributes about mangrove distribution and 12 types of reclamation pressure. Sensitivity is a dose-response of mangrove system to reclamation, it is an interaction between reclamation and mangrove conditions. Sensitivity, as an interlinked factor between the other two aspects, was calculated with potential loss and potential structure change of mangroves to different reclamation pressures. Resilience refers to recovery capacities of mangrove system to reclamation activities, resilience was represented by both two internal indicators (species numbers and ratio of non-threaten species) and external resilience indicator (ecological management).

Vulnerability of mangroves can be characterized by degree to which mangrove is likely to experience damage during coastal development. Vulnerability of mangroves is predicated on trade-offs between economic development and coastal ecosystem services maintenance, as they are affected by interconnections operating at different spatiotemporal and functional scales. Based on vulnerability framework, the Habitat Risk Assessment model in InVEST was used to calculate spatial vulnerability of mangroves in each province while adapting to coastal reclamation (Sharp et al., 2015). Combination of vulnerability framework and InVEST model produces spatial qualitative estimate of potential risks in terms of vulnerability value while exposed to reclamation, which differentiates areas with relatively high or low exposure to reclamation threats.

## 4 Results

### 4.1 General distribution of mangroves

In total, 25,829 ha mangroves were identified in five coastal provinces of mainland China in 2015 (Fig. 5), 96.14% of which were distributed in southern part of coastal provinces (Hainan, Guangxi and Guangdong Province), mainly located in the Beibu Gulf Economic Rim (the economic region surrounding around China’s southwestern coastal area). About 11,115 ha (43%) mangroves was classified in Guangdong Province, where many coastal areas are dominated by sedimentary environments. Followed by Guangxi Province with a number of 9,297 ha mangroves, the third largest area of mangroves was 4,420 ha in Hainan Province. About 963 ha and 34 ha mangroves were detected in Fujian Province and Zhejiang Province respectively.

**Fig. 5.**
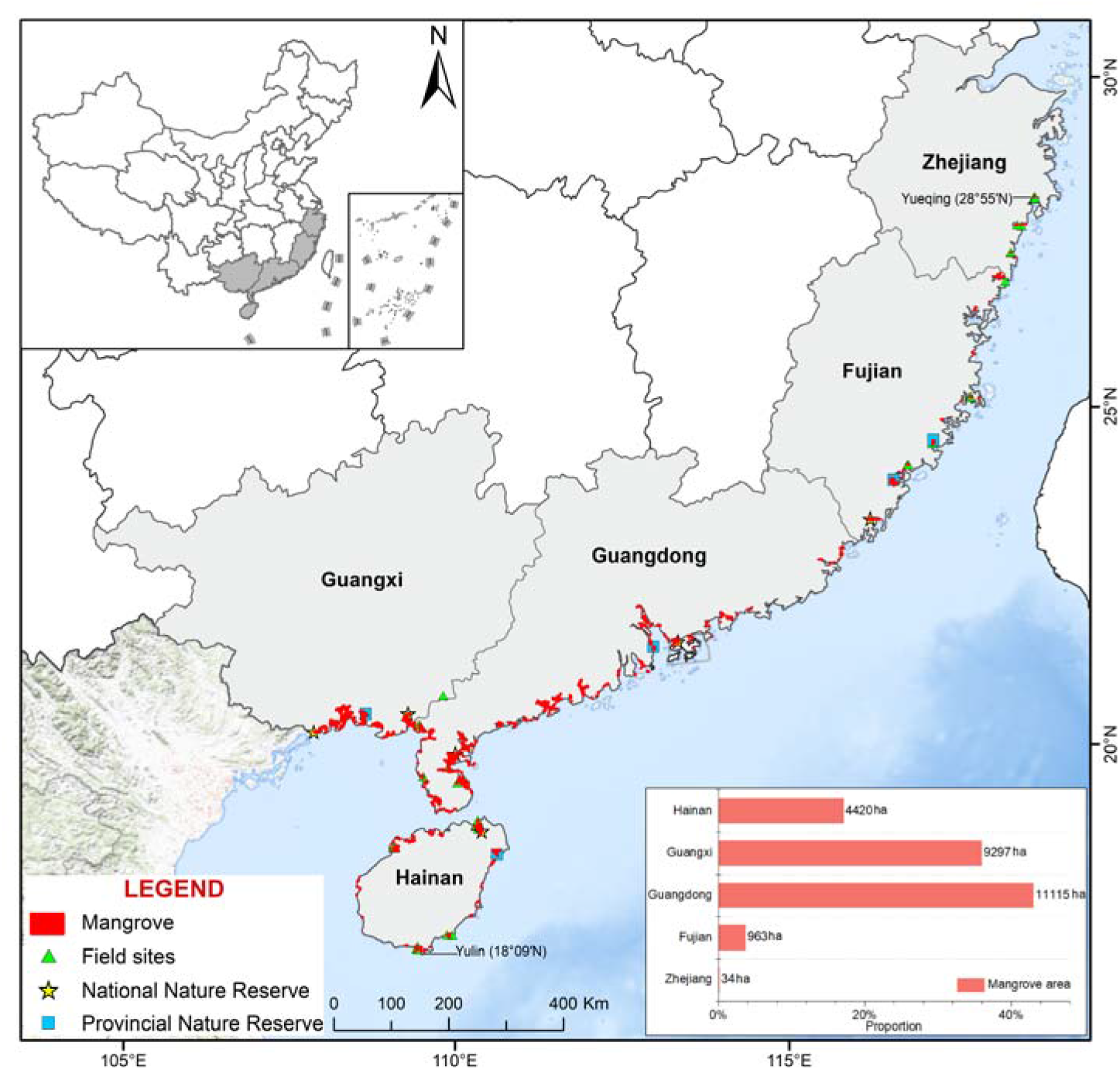
Mangroves distribution in five coastal provinces.

### 4.2 Principal disturbance agents to mangroves

Landscape conversion from natural coastline to artificial coastline was the key driver of mangrove degradation. By overlapping coastline and mangroves, we found that only 7% of mangrove was adjacent to natural coastline types (Fig. 6). Most of mangroves were exposed to anthropogenic disturbances caused by redeveloped coastlines, where 64% of mangrove was located near aquaculture ponds and 29% of mangrove was threatened by filled land. It suggests that 93% of mangrove was squeezed by human-built bank and dam toward ocean, and they were threatened by floods on the marine side.

**Fig. 6.**
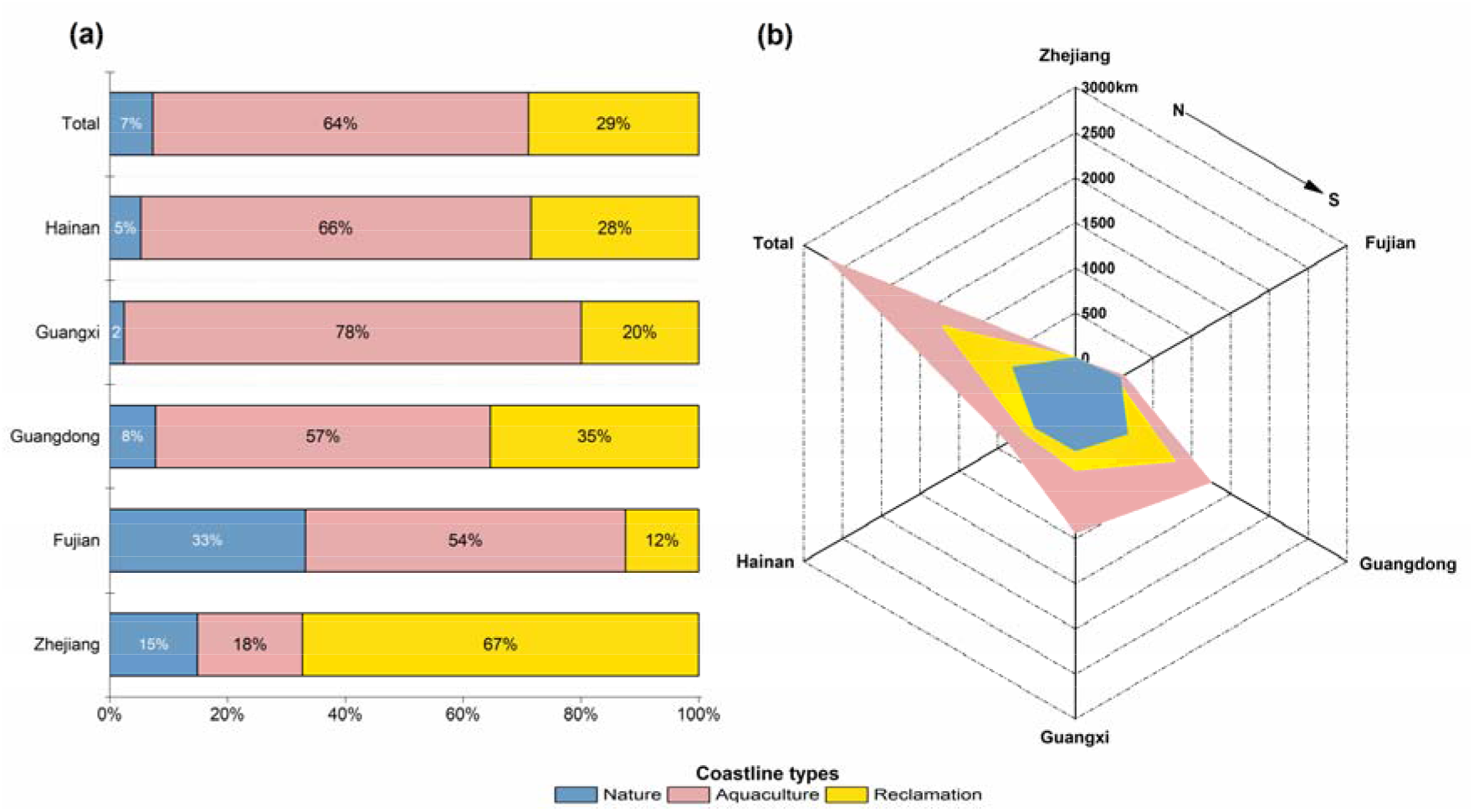
Proportions (a) and lengths (b) of different coastline types in 2015.

In a local scale, reclamation activities in Zhejiang Province resulted in the highest density of filled coastline among all coastal provinces. Followed by Guangxi Province, high percentage (98%) of artificial coastline was adjacent to mangrove, due to large areas of aquaculture feeding ponds. In all these five provinces, the highest ratio (30%) of natural coastline adjacent to mangroves appeared in Fujian Province. Even though the coverage of aquaculture in Fujian, Guangdong and Hainan Province were lower than that in Guangxi, but aquaculture land in all provinces covered more than that of natural land. Guangdong was the province which has the longest coastline both artificial and nature, followed by Guangxi, demonstrating that these two provinces need to be given priority (Fig. 6b).

### 4.3 Spatial distribution of mangrove vulnerability

In this study, spatial distribution of vulnerability in mangroves was derived by combining vulnerability framework and Habitat Risk Assessment (HAR) model in InVEST (Fig. 7). We found that 29% of mangrove areas was identified under high vulnerability, only 24% of them was estimated as area with low vulnerability. The average value of vulnerability in all five provinces were ranked from high to low as Zhejiang, Hainan, Guangxi, Guangdong, and Fujian. As can be seen from Fig. 8, all mangroves in Zhejiang Province were under high vulnerability, which indicates high risk of degradation in this region. Mangroves in Zhejiang Province were planted species, due to its climate conditions and high coverage of artificial coastline, there is limited suitable natural coastal areas for mangroves. About 64.41% of mangroves in Hainan Province were observed under high vulnerability, only 9.44% of them were remained in low human influences. The main reason of such high vulnerability in this region was caused by aquaculture pollution and high ratio of endangered species. Nearly 25% of mangroves in both Guangxi and Fujian Province was classified with high vulnerability, it was because of marine aquaculture and coastal redevelopment, all of which have led to significant degradation and ecological disturbances to mangroves.

**Fig. 7.**
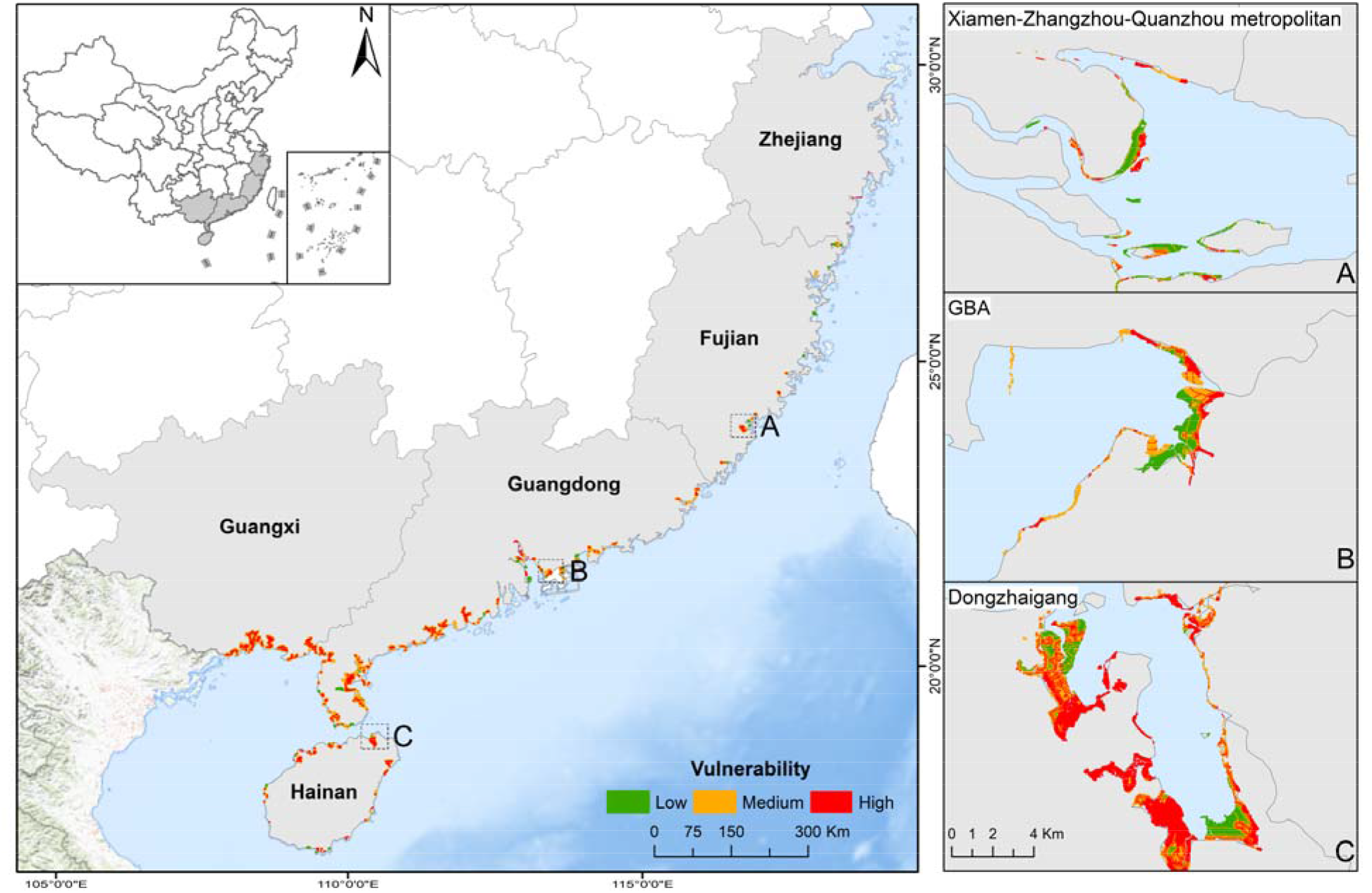
Distribution of mangrove vulnerability to anthropogenic disturbances along China’s coastal area. (GBA: Guangdong-Hongkong-Macao Greater Bay Area).

**Fig. 8.**
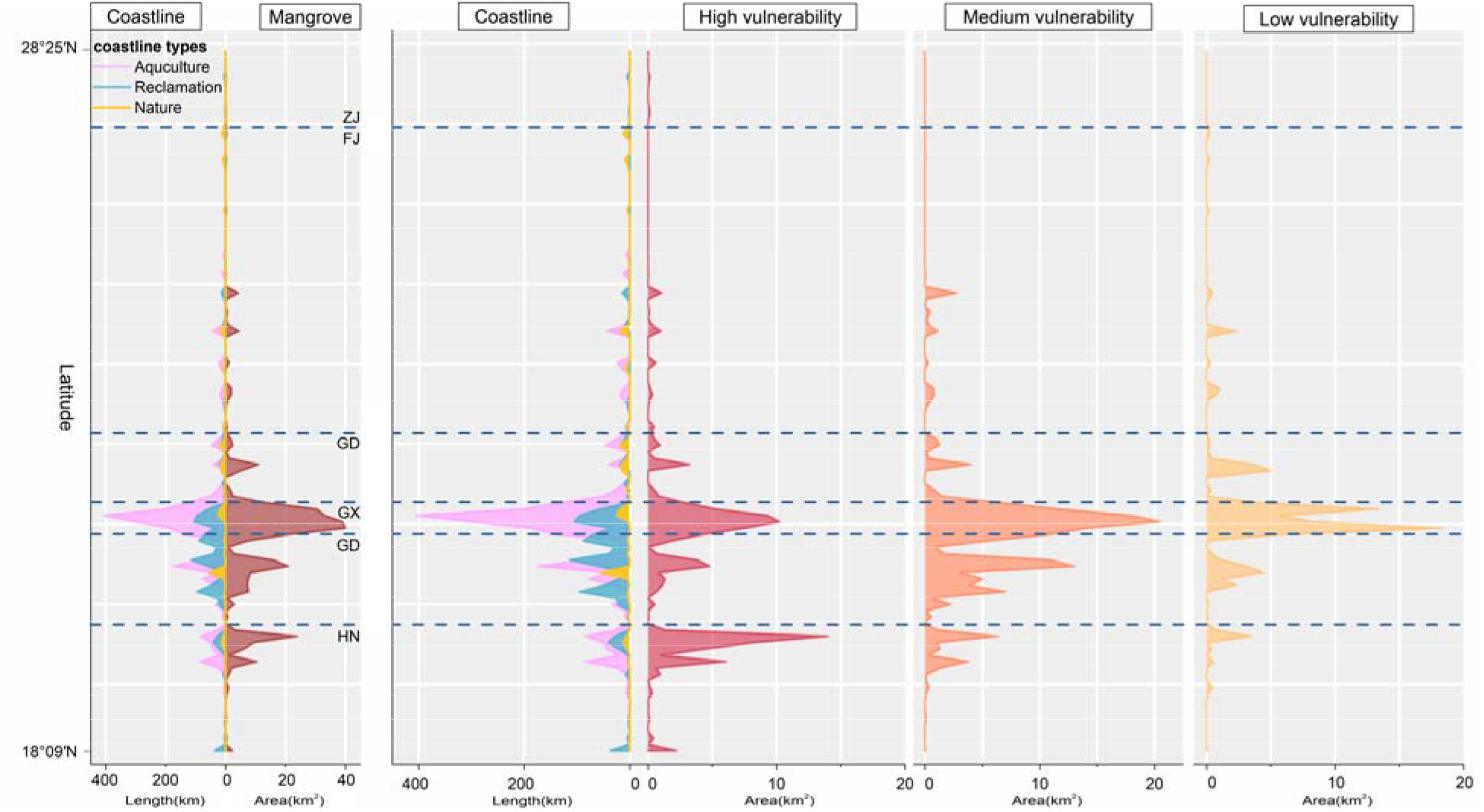
Spatial distribution of coastline with mangrove and different level of vulnerability. The left side of Y axis shows the latitude of different types of coastline (aquaculture, reclamation, and natural) from Zhejiang Province to Hainan Province (ZJ: Zhejiang Province; FJ: Fujian Province; GD: Guangdong Province; GX: Guangxi Province; HN: Hainan Province), the right side of Y axis represents area of mangroves and different level of vulnerability spatially. The left X axis is the length of mangroves and right X axis is the area of different level of vulnerability.

Among all these selected vulnerability indicators, aquaculture and reclamation coastlines were the key drivers of mangroves degradation. Areas with high vulnerability in Guangxi and Hainan Province was found near large areas of aquaculture and reclamation. The overall distribution of high, medium and low vulnerability was following similar trend (Fig. 8), the largest groups of vulnerability (peak values) appeared in Guangxi, Guangdong and Hainan Province. However, peak values of three levels of vulnerability were slightly different, while that of medium and low vulnerability were in Guangxi Province, and mangroves in Hainan had the largest area of high vulnerability. In general, based on the distribution of coastline and vulnerability, we found that aquaculture was primarily threats to high risk of mangroves degradation in these five provinces.

In a local scale, mangroves in urban agglomerations experienced higher vulnerability than that rest of mangrove (Fig. 7). As the Guangdong-Hong Kong-Macao Greater Bay Area (GBA) becomes an important economic zone in China, rapid urbanization in coastal areas lead to high level of disturbance caused by human activities, which is the principal threat to local mangrove ecosystem. In the meantime, Guangdong Province has the largest area of mangrove in all five provinces, which includes 7 natural protected areas for mangrove conservation. These protected areas help to maintain key functions of local mangrove ecosystems under coastal reclamation. The northern boundary of natural mangroves is located in Fujian Province, the Xiamen-Zhangzhou-Quanzhou metropolitan in this Province was also exposed to relative high vulnerability due to extensive reclamation activities. Even if protected in a national nature reserve, mangroves in Dongzhaigang also faced serious threats due to the substantially increased aquaculture ponds and construction area.

### 4.4 Accuracy assessment

Our results show that the map of mangrove forests in 2015 showed a high user’s accuracy of 97.4% and producer’s accuracy of 82.2%. We also compared our results with similar research in China to quantify the reliability of our interpretation (Jia et al., 2018). Detected total area of mangroves (25,829 ha) in our research was comparable with their results derived from Landsat archive (22,419 ha). Due to the coarse resolution of Landsat archive (30 m), small mangrove fragments in urban agglomeration identified in our study was not able to be classified by using Landsat images. Therefore, the identified area of mangrove in our study is larger than their result. The main difference between these two datasets was in the Pearl River Estuary and Xiamen-Zhangzhou-Quanzhou metropolitan. In order to improve our accuracy of mangrove identification in those regions, field investigation was carried out in summer time. Overall, the validation results suggest that our mangrove map is mainly consistent with ground truth and similar research in China.

## 5 Discussion

### 5.1 Impacts of principal threats on vulnerability distribution to local ecosystems

Due to continuous influences of coastal reclamation and aquaculture farming, mangroves are exposed to dynamic and extensive threats in local region. Conversion of mangroves to aquaculture ponds was encouraged by local governments, which becomes main income of coastal regions. Among all types of coastal development activity in terms of land use change, the most significant replacement of land use was aquaculture (64% of total area), which leads to high vulnerability of local mangroves.

As comparing all three levels of vulnerability distribution, we found that mangroves with medium level of vulnerability was the most sensitive area to aquaculture factor. The overall distribution pattern of medium vulnerability was coordinated with that of aquaculture. Except in Zhejiang Province, aquaculture was the domain land use type in local mangrove areas, which threatened more than half of mangroves in our study area. Threat of aquaculture in driving mangrove deforestation is not only a problem in China, but also was found in southeast Asia countries, such as Thailand, Indonesia, Vietnam, and the Philippines (Primavera, 2000; Richards & Friess, 2016).

Land redevelopment and infrastructure construction was another direct factor leading to mangrove loss. Even it influences on vulnerability was not that significant as aquaculture, it shows high correlations with high level of vulnerability, especially in the areas dominated by reclamation coastlines in Hainan and Fujian Province. In those areas adjacent to long artificial coastline, it also addresses high level of influence on vulnerability. Extensive and large area of land reclamation occurred in Pearl River Delta (Guangdong Province) and Beibu Gulf (Guangxi Province), these regions were shown with high vulnerability of mangrove degradation. Coastal artificial infrastructures, such as large industrial complexes, coastal interstate highways, and airports, occupied areas suitable for mangrove growth, and accounted for 29% of total mangrove areas in mainland China. Local government historically aimed to increase terrestrial land through expansion towards ocean, especially in the most developed metropolises, such as GBA, Bohai Bay (Tianjin Province), Hangzhou Bay (Zhejiang Province) and Min River estuary (Fujian Province). High intensity of human disturbances toward ocean poses potential stress to mangrove ecosystem, which reduce the buffer zone of mangroves in adapting to sea level rise.

### 5.2 Threats to local ecosystem services: habitats and biodiversity

Mangroves degradation and loss is not only bounded with mangrove ecosystem itself, furthermore, spillover effects of mangroves degradation have significant influences on ecosystem processes and services that they provide. Vulnerability of habitats and biodiversity increased as structure and functionality of local ecosystems were challenged by anthropogenic disturbances. Within areas of high vulnerability in mangroves, the most challenges to local ecosystems were habitats degradation and biodiversity loss (Arkema et al., 2014). Nearly 35 ha of mangrove disappeared in Futian mangrove conservation area because of city construction, which leads to 39% of bird density loss and 45% of species reduction (Wang & Wang, 2007). Species extinctions can be followed by loss in functional diversity, particularly in mangroves with high vulnerability. Further degradation in mangroves is likely to be followed by accelerated functional losses. New top-down policies adopted by China’s central government in 2016 has imposed a conception of ‘Redline’, a boundary delineating a coastal strip in which to preserve existing natural habitats, both for maintaining ecosystem function and to reduce conflicts between natural processes (exacerbated by accelerated sea-level rise) and human settlements and infrastructure. The Central Leading Group for Comprehensively Deepening Reforms of China approved a series of decisions of Redline policy in 2016. It comprises “Opinions on Delimiting and Guarding Ecological Protection Redline”, “Program for Wetland Protection and Restoration”, and “Rules on Coastal Line Protection and Utilization”, which all emphasize the integration of “One Redline Map” in China’s coastal area. China’s coastal Redline would be challenged by 2020 in maintaining 35% (6,300 km) of coastal line, which is the goal set by the central government in protecting coastal line (Larson, 2015). Consequently, transitioning the coastline from natural to artificial, through such methods as land claim or armoring, makes coastal areas more vulnerable in terms of lost biodiversity and increased natural hazards (Arkema et al., 2015; Steffen et al., 2015). Mangroves, as a typical coastal ecosystem included in the Redline area, maintain key functions of local ecosystems, they also contribute to biodiversity conservation.

In 2018, China’s governments announced to halt all coastal land reclamation that related to business land use. But degradation of mangroves and irreversible landscape changes are still challenging in maintain key functions of coastal ecosystems. As mangrove habitats are smaller or fragmented, their essential ecosystem services may be lost within and beyond its boundary. The central government should act quickly to nominate and protect vulnerable coastal sites, because this would be consistent with the national Redline program and reclamation restoration. The local governments of coastal provinces and cities should also proactively adopt effective measures of their own, consistent with the national Redline policies, to protect vulnerable coastal ecosystem.

### 5.3 Limitations

There are some uncertainties in this study. The 900-m-buffer of coastal line and the 300 m interval in the buffer zone were chosen based on general distribution of mangroves in our study area, which may different while implying to other regions. Moreover, in the current analysis, all 8 indicators in vulnerability assessment were treated with equal weight, which was set according to current states of coastal region in China. This may various due to spatial heterogeneity in different area, and need to adjust while applying to other cases.

Spatial outputs of mangrove vulnerability in this research provide a qualitative representation of the relative contributions of anthropogenic disturbances to coastal vulnerability and highlight the potential threats of reclamation in degrading mangrove ecosystems. Further information on social, economic and coastal disaster are helpful in giving a comprehensive assessment of mangroves ecosystem and better informing development strategies and permitting in setting conservation priorities, monitoring deforestation and forest degradation.

## 6 Conclusion

Mangroves are under threats from different types of anthropogenic disturbances, coastal rapid development, in terms of aquaculture and reclamation, has been a principal driver of mangrove loss and degradation in China, and it becomes a growing concern worldwide. These two factors were the main drivers of medium and high level of vulnerability. The continuously altered, destroyed or transformed mangroves with high vulnerability is an early warning signal of dramatic functionality loss of such major coastal ecosystem, which initiates more research of mangrove in coastal urban environments.

Mangrove areas of high vulnerability suggest relative high level of combinate anthropogenic disturbances from conversion for alternative uses (aquaculture, urban construction), which can be prioritized for the development of conservation strategies. Distribution of mangrove vulnerability also located the hotspots of key functionality loss and the vulnerable urban areas with high risk of storm surges or climate-related disasters. As we estimated, the major disturbance agent to mangroves was aquaculture ponds, especially in Guangxi and Hainan Provinces. These findings indicate an urgent need for implementing adaptive strategies to serve mangroves from anthropogenic disturbances, and provides a spatially-explicit assessment for navigating mangrove conservation.

## Acknowledgements

We would like to appreciate Professor Wenqing Wang at Xiamen University who provided valuable comments and suggestions. We also thank Dr. Diego Rybski in Potsdam Institute for Climate Impact Research for his insightful comments and suggestions. Financial supports were provided by the Programs of Science and Technology on Basic Resources Survey for the Ministry of Science and Technology of China (No. 2017FY100701), the National Natural Science Foundation of China (NSFC) Grants (No. 41976208 and 41701205), and the Fundamental Research Funds for the Central Universities of China (No. 20720190089).

## Notes

### Competing Interest Statement

The authors have declared no competing interest.

## References

Adger, W. N. (2006). Vulnerability. Global Environmental Change, 16(3), 268–281. https://doi.org/10.1016/j.gloenvcha.2006.02.006

Arkema, K. K., Verutes, G., Bernhardt, J. R., Clarke, C., Rosado, S., Canto, M., et al. (2014). Assessing habitat risk from human activities to inform coastal and marine spatial planning: a demonstration in Belize. Environmental Research Letters, 9(11), 114016. https://doi.org/10.1088/1748-9326/9/11/114016

Arkema, K. K., Verutes, G. M., Wood, S. A., Clarke-Samuels, C., Rosado, S., Canto, M., et al. (2015). Embedding ecosystem services in coastal planning leads to better outcomes for people and nature. Proceedings of the National Academy of Sciences, 112(24), 7390–7395. https://doi.org/10.1073/pnas.1406483112

Bell, J., & Lovelock, C. E. (2013). Insuring Mangrove Forests for Their Role in Mitigating Coastal Erosion and Storm-Surge: An Australian Case Study. Wetlands, 33(2), 279–289. https://doi.org/10.1007/s13157-013-0382-4

Branoff, B. L. (2017). Quantifying the influence of urban land use on mangrove biology and ecology: A meta-analysis. Global Ecology and Biogeography, 26(11), 1339–1356. https://doi.org/10.1111/geb.12638

Crooks, S., Sutton-Grier, A. E., Troxler, T. G., Herold, N., Bernal, B., Schile-Beers, L., & Wirth, T. (2018). Coastal wetland management as a contribution to the US National Greenhouse Gas Inventory. Nature Climate Change, 8(12), 1109–1112. https://doi.org/10.1038/s41558-018-0345-0

DasGupta, R., & Shaw, R. (2013). Changing perspectives of mangrove management in India-An analytical overview. Ocean & Coastal Management, 80, 107–118. https://doi.org/10.1016/j.ocecoaman.2013.04.010

Doody, J. P. (2013). Coastal squeeze and managed realignment in southeast England, does it tell us anything about the future? Ocean & Coastal Management, 79(Supplement C), 34–41. https://doi.org/10.1016/j.ocecoaman.2012.05.008

Duke, N., Nagelkerken, I., Agardy, T., Wells, S., & Van Lavieren, H. (2014). The importance of mangroves to people: A call to action. United Nations Environment Programme World Conservation Monitoring Centre (UNEP-WCMC), Cambridge.

Elmqvist, T., Fragkias, M., Goodness, J., Güneralp, B., Marcotullio, P. J., McDonald, R. I., et al. (2013). Urbanization, Biodiversity and Ecosystem Services: Challenges and Opportunities. Dordrecht: Springer Netherlands. https://doi.org/10.1007/978-94-007-7088-1

Estoque, R. C., Myint, S. W., Wang, C., Ishtiaque, A., Aung, T. T., Emerton, L., et al. (2018). Assessing environmental impacts and change in Myanmar’s mangrove ecosystem service value due to deforestation (2000-2014). Global Change Biology, 24(11), 5391–5410. https://doi.org/10.1111/gcb.14409

Flores-de-Santiago, F., Serrano, D., Flores-Verdugo, F., & Monroy-Torres, M. (2017). Application of a simple and effective method for mangrove afforestation in semiarid regions combining nonlinear models and constructed platforms. Ecological Engineering, 103, 244–255. https://doi.org/10.1016/j.ecoleng.2017.04.008

Friess, D. A., & Webb, E. L. (2014). Variability in mangrove change estimates and implications for the assessment of ecosystem service provision. Global Ecology and Biogeography, 23. https://doi.org/10.1111/geb.12140

Friess, D. A., Thompson, B. S., Brown, B., Amir, A. A., Cameron, C., Koldewey, H. J., et al. (2016). Policy challenges and approaches for the conservation of mangrove forests in Southeast Asia: Conservation of Mangrove Forests. Conservation Biology, 30(5), 933–949. https://doi.org/10.1111/cobi.12784

Hamilton, S. E., & Casey, D. (2016). Creation of a high spatio-temporal resolution global database of continuous mangrove forest cover for the 21st century (CGMFC-21): CGMFC-21. Global Ecology and Biogeography, 25(6), 729–738. https://doi.org/10.1111/geb.12449

Hamilton, S. E., & Friess, D. A. (2018). Global carbon stocks and potential emissions due to mangrove deforestation from 2000 to 2012. Nature Climate Change, 8(3), 240–244. https://doi.org/10.1038/s41558-018-0090-4

Intergovernmental Panel on Climate Change (IPCC). (2014). IPCC Fifth Assessment Report (AR5): Climate Change 2014: Impacts, Adaptation, and Vulnerability. Part A: Global and Sectoral Aspects. Contributionof Working Group II to the Fifth Assessment Report of the IntergovernmentalPanel on Climate Change. (p. 169). Cambridge University Press, Cambridge, UK.

Jia, M., Wang, Z., Zhang, Y., Mao, D., & Wang, C. (2018). Monitoring loss and recovery of mangrove forests during 42 years: The achievements of mangrove conservation in China. International Journal of Applied Earth Observation and Geoinformation, 73, 535–545. https://doi.org/10.1016/j.jag.2018.07.025

Koh, C., & Khim, J. S. (2014). The Korean tidal flat of the Yellow Sea: Physical setting, ecosystem and management. Ocean & Coastal Management, 102, 398–414. https://doi.org/10.1016/j.ocecoaman.2014.07.008

Larson, C. (2015). Migratory bird populations in Asia are crashing as Yellow Sea habitat dwindles. Science, 350(6257), 150–152.

Lovelock, C. E., Cahoon, D. R., Friess, D. A., Guntenspergen, G. R., Krauss, K. W., Reef, R., et al. (2015). The vulnerability of Indo-Pacific mangrove forests to sea-level rise. Nature, 526(7574), 559–563. https://doi.org/10.1038/nature15538

McLeod, E., & Salm, R. V. (2006). Managing mangroves for resilience to climate change. The World Conservation Union (IUCN), Gland, Switzerland.

Mcleod, E., Chmura, G. L., Bouillon, S., Salm, R., Björk, M., Duarte, C. M., et al. (2011). A blueprint for blue carbon: toward an improved understanding of the role of vegetated coastal habitats in sequestering CO _2_. Frontiers in Ecology and the Environment, 9(10), 552–560. https://doi.org/10.1890/110004

Millennium Ecosystem Assessment. (2005). Ecosystems and human well-being: synthesis. Washington, DC: Island Press.

Moran, E. F., Lopez, M. C., Moore, N., Müller, N., & Hyndman, D. W. (2018). Sustainable hydropower in the 21st century. Proceedings of the National Academy of Sciences, 115(47), 11891–11898. https://doi.org/10.1073/pnas.1809426115

Mukherjee, N., Sutherland, W. J., Khan, M. N. I., Berger, U., Schmitz, N., Dahdouh-Guebas, F., & Koedam, N. (2014). Using expert knowledge and modeling to define mangrove composition, functioning, and threats and estimate time frame for recovery. Ecology and Evolution. https://doi.org/10.1002/ece3.1085

Murray, N. J., Phinn, S. R., DeWitt, M., Ferrari, R., Johnston, R., Lyons, M. B., et al. (2018). The global distribution and trajectory of tidal flats. Nature. https://doi.org/10.1038/s41586-018-0805-8

Nandy, S., & Kushwaha, S. P. S. (2011). Study on the utility of IRS 1D LISS-III data and the classification techniques for mapping of Sunderban mangroves. Journal of Coastal Conservation, 15(1), 123–137. https://doi.org/10.1007/s11852-010-0126-z

Nortey, D. D. N., Aheto, D. W., Blay., Jonah, F. E., & Asare, N. K. (2016). Comparative Assessment of Mangrove Biomass and Fish Assemblages in an Urban and Rural Mangrove Wetlands in Ghana. Wetlands, 36(4), 717–730. https://doi.org/10.1007/s13157-016-0783-2

Primavera, J. H. (2000). Development and conservation of Philippine mangroves: institutional issues. Ecological Economics, 35(1), 91–106. https://doi.org/10.1016/S0921-8009(00)00170-1

Ramesh, R., Chen, Z., Cummins, V., Day, J., D’Elia, C., Dennison, B., et al. (2015). Land-Ocean Interactions in the Coastal Zone: Past, present & future. Anthropocene, 12, 85–98. https://doi.org/10.1016/j.ancene.2016.01.005

Richards, D. R., & Friess, D. A. (2016). Rates and drivers of mangrove deforestation in Southeast Asia, 2000-2012. Proceedings of the National Academy of Sciences, 113(2), 344–349. https://doi.org/10.1073/pnas.1510272113

Rivera-Monroy, V., Lee, S. Y., Kristensen, E., & Twilley, R. (2017). Mangrove ecosystems: a global biogeographic perspective. New York, NY: Springer International Publishing.

Rocha, J. C., Peterson, G., Bodin, Ö., & Levin, S. (2018). Cascading regime shifts within and across scales, 31.

Romañach, S. S., DeAngelis, D. L., Koh, H. L., Li, Y., Teh, S. Y., Raja Barizan, R. S., & Zhai, L. (2018). Conservation and restoration of mangroves: Global status, perspectives, and prognosis. Ocean & Coastal Management, 154, 72–82. https://doi.org/10.1016/j.ocecoaman.2018.01.009

Sajjad, M., Li, Y., Tang, Z., Cao, L., & Liu, X. (2018). Assessing Hazard Vulnerability, Habitat Conservation, and Restoration for the Enhancement of Mainland China’s Coastal Resilience. Earth’s Future, 6(3), 326–338. https://doi.org/10.1002/2017EF000676

Seto, K. C., Güneralp, B., & Hutyra, L. R. (2012). Global forecasts of urban expansion to 2030 and direct impacts on biodiversity and carbon pools. Proceedings of the National Academy of Sciences, 109(40), 16083–16088. https://doi.org/10.1073/pnas.1211658109

Sharp, R., Tallis, H. T., Ricketts, T., Guerry, A. D., Wood, S. A., Chaplin-Kramer, R., et al. (2015). InVEST +VERSION+ User’s Guide. The Natural Capital Project, Stanford University, University of Minnesota, The Nature Conservancy, and World Wildlife Fund. Retrieved from http://data.naturalcapitalproject.org/nightly-build/release_default/release_default/documentation/

Shi, H., Chen, J., Liu, S., & Sivakumar, B. (2019). The role of large dams in promoting economic development under the pressure of population growth. Sustainability, 11(10), 2965. https://doi.org/10.3390/su11102965

Silliman, B. R., Bertness, M. D., & Grosholz, E. D. (2009). Human Impacts on Salt Marshes: A Global Perspective (First edition). Berkeley: University of California Press.

Spalding, M. (2010). World Atlas of Mangroves. London, UK: EarthScan.

State Oceanic Administration, People’s republic of China. (2016). Report on the management of the use of sea areas (2015). State Oceanic Administration, People’s republic of China, Beijing, China. Retrieved from http://www.soa.gov.cn/zwgk/hygb/hysyglgb/201604/t20160429_51327.html

Steffen, W., Richardson, K., Rockström, J., Cornell, S. E., Fetzer, I., Bennett, E. M., et al. (2015). Planetary boundaries: Guiding human development on a changing planet. Science, 347(6223). https://doi.org/10.1126/science.1259855

Sutton-Grier A E., Wowk K., Bamford H. (2015). Future of our coasts: The potential for natural and hybrid infrastructure to enhance the resilience of our coastal communities, economies and ecosystems [J]. Environmental Science and Policy, 51, 137–148. https://doi.org/10.1016/j.envsci.2015.04.006

Tian, B., Wu, W., Yang, Z., & Zhou, Y. (2016). Drivers, trends, and potential impacts of long-term coastal reclamation in China from 1985 to 2010. Estuarine, Coastal and Shelf Science, 170, 83–90. https://doi.org/10.1016/j.ecss.2016.01.006

Turner II, B. L., Kasperson, R. E., Matson, P. A., McCarthy, J. J., Corell, R. W., Christensen, L., et al. (2003). A framework for vulnerability analysis in sustainability science. Proceedings of the National Academy of Sciences, 100(14), 8074–8079. https://doi.org/10.1073/pnas.1231335100

Valiela, I., Bowen, J. L., & York, J. K. (2001). Mangrove Forests: One of the World’s Threatened Major Tropical Environments. BioScience, 51(10), 807. https://doi.org/10.1641/0006-3568(2001)051[0807:MFOOTW]2.0.CO;2

Ventura, A. de O. B., & Lana, P. da C. (2014). A new empirical index for assessing the vulnerability of peri-urban mangroves. Journal of Environmental Management, 145, 289–298. https://doi.org/10.1016/j.jenvman.2014.04.036

Wang, Q., Li, Y., & Li, Y. (2018). Realizing a new resilience paradigm on the basis of land-water-biodiversity nexus in a coastal city. Ocean & Coastal Management. https://doi.org/10.1016/j.ocecoaman.2018.09.004

Wang, W., & Wang, M. (2007). Mangroves in China (in Chinese). China Science Press.

Webb, E. L., Friess, D. A., Krauss, K. W., Cahoon, D. R., Guntenspergen, G. R., & Phelps, J. (2013). A global standard for monitoring coastal wetland vulnerability to accelerated sea-level rise. Nature Climate Change, 3(5), 458–465. https://doi.org/10.1038/nclimate1756

White, G. F. (1974). Natural hazards: Local, national, global. Oxford University Press, New York.

Xin, K., Huang, X., Hu, J., Li, C., Yang, X., & Arndt, S. K. (2014). Land use Change Impacts on Heavy Metal Sedimentation in Mangrove Wetlands—A Case Study in Dongzhai Harbor of Hainan, China. Wetlands, 34(1), 1–8. https://doi.org/10.1007/s13157-013-0472-3

Xu, Y., Shen, Z.-H., Ying, L.-X., Ciais, P., Liu, H.-Y., Piao, S., et al. (2016). The exposure, sensitivity and vulnerability of natural vegetation in China to climate thermal variability (1901–2013): An indicator-based approach. Ecological Indicators, 63, 258–272. https://doi.org/10.1016/j.ecolind.2015.12.023

Yim, J., Kwon, B.-O., Nam, J., Hwang, J. H., Choi, K., & Khim, J. S. (2018). Analysis of forty years long changes in coastal land use and land cover of the Yellow Sea: The gains or losses in ecosystem services. Environmental Pollution, 241, 74–84. https://doi.org/10.1016/j.envpol.2018.05.058

